# *Coro2a*, An Expression Quantitative Trait Gene Underlying *Estq1*, Controls Uterine Responsiveness to Estradiol

**DOI:** 10.1101/2021.02.28.433254

**Authors:** Joseph Findley, Elizabeth McGee, Chin-Yo Lin, Dimitry Krementsov, Cory Teuscher

## Abstract

The response of the uterus to 17β-estradiol (E_2_) is genetically controlled, with marked variation observed depending on the mouse strain studied. Previous studies from our laboratory using high (C57BL6/J; B6) and low uterine responders (C3H/HeJ; C3H) to E_2_ led to the identification of *Estq1-Estq5*, five quantitative trait loci (QTL) controlling phenotypic variation in uterine growth and eosinophilic leukocyte infiltration. Transcriptional profiling subsequently identified coronin actin binding protein 2A (*Coro2a*) as a candidate for an expression quantitative trait gene (eQTG) underlying *Estq1*. Here we show that like mRNA expression, allele specific expression of CORO2A in both the uterine epithelium and stroma similarly co-segregates with uterine responsiveness. Utilizing B6.C3H congenic mice carrying the *Coro2a^C3H^* allele, we confirm the genetic linkage between *Coro2a* and *Estq1*. Lastly, we show that B6-*Coro2a^-/-^* and B6-*Coro2a^B6/-^* mice phenocopy uterine responsiveness of mice carrying the *Coro2a^C3H/C3H^* and *Coro2a^B6/C3H^* alleles respectively, consistent with a haploinsufficiency model of genetic control. Together, these results identify *Coro2a* as the eQTG underlying *Estq1*, and establish a critical role for *Coro2a* in E_2_-regulated responses.

## Introduction

Estrogens are female sex hormones that regulate a variety of physiological processes including reproductive development and function, wound healing, and bone growth (1, 2). Estrogens are primarily known for their role in reproductive tissues, particularly the ability of 17β-estradiol (E_2_), the primary estrogen secreted by the ovary, to induce cellular proliferation and growth of the uterus. E_2_ elicits a marked increase in uterine epithelial cell proliferation, vascular permeability, and water imbibition, followed by an influx of leukocytes into the uterine stroma. Together, these comprise the classical uterotrophic response (3). In addition to regulating normal physiological states, estrogens have a well-established role in determining susceptibility to disease, particularly cancer, in reproductive tissues (4–7).

Early studies in mice demonstrated that the E_2_-regulated uterotrophic response is genetically controlled, with marked variation in tissue growth and (or) regression depending on the strain of mouse studied (8, 9). More recently, our laboratory has demonstrated that the infiltration of leukocytes, particularly eosinophils, into the uterine stroma is genetically determined (10). Genetic studies using inbred strains of mice that are uterine high-responders (C57BL6/J; B6) or low-responders (C3H/HeJ; C3H) to E_2_, resulted in the identification of five interacting quantitative trait loci (QTL) that control phenotypic variation in uterine growth and leukocyte infiltration *(Estq1-Estq5)* (10, 11).

Subsequently, utilizing genome-wide transcriptional profiling and RT qPCR, we identified coronin actin binding protein 2A (*Coro2a*/CORO2A) as a potential positional candidate for an expression quantitative trait gene (eQTG) underlying *Estq1* (12). Specifically, under basal conditions *Coro2a* mRNA expression was five-fold greater in the uteri of high-responder B6 mice compared to low-responder C3H mice, and intermediate in (B6 × C3H) F_1_ hybrids (B6C3). These findings indicate that *Coro2a*, acting as an eQTG would be doing so during the acute/rapid phase (0-1 hr) of the early uterotophic response (13).

CORO2A, also known as CLIPINB, IR10, or WDR2, is a member of the coronin family of actin-binding proteins (14) and is a component of the nuclear receptor co-repressor (NCoR) complex (15, 16) that mediates toll-like receptor-induced NCoR turnover by interacting with oligomeric nuclear actin; thereby, leading to derepression of inflammatory response genes (17). This mechanism may play a similar role in E_2_-regulated responses given that NCoR interacts with estrogen receptor-1 (ESR1) and estrogen receptor-2 (ESR2), and that E_2_-induced NCoR turnover occurs within the first 10 min following E_2_ treatment of MCF-7 cells (18–22). Moreover, *CORO2A* is overexpressed in breast cancer (23, 24), a malignancy predominantly associated with aberrant estrogen signaling, and knockdown of CORO2A expression disrupts cell proliferation and colony formation of estrogen-dependent MCF-1 breast cancer cells (25).

Consequently, the hypothesis that we tested in this study is that *Coro2a* is the eQTG underlying *Estq1*. Given that low- and high-uterine responsiveness to E_2_ in C3H and B6 mice co-segregates with differential *Coro2a*/CORO2A expression, and haploinsufficiency is seen in B6C3 mice, we utilized deficiency-complementation testing (26, 27) to determine if *Coro2a* underlies *Estq1*. We show that B6-*Coro2a*^-/-^ and B6-Coro2a^B6/-^ mice phenocopy the low uterine responsiveness of C3H mice, and haploinsufficiency in B6C3 mice, respectively. Taken together, these results identify *Coro2a* as the eQTG underlying *Estq1*, and provide genetic evidence for a potential role for *Coro2a* in E_2_-regulated responses.

## Materials and Methods

### Mice

Female C57BL/6J (B6) and C3H/HeJ (C3H/C3) mice were purchased from the Jackson Laboratory (Bar Harbor, ME) and acclimatized for two weeks prior to experimentation. B6-^*Coro2atm1(KOMP)Wtsi*^ (B6-*Coro2a^-/-^*) mice were provided by Dr. Christopher K. Glass (University of California, San Diego, CA). *B6.C3H-Bbaa1a* (11.6-77.8 Mbp) mice were obtained from Dr. Janis J. Weis (University of Utah, Salt Lake City, UT) (28, 29). All animals, including (B6 × C3H) and (B6 × B6-*Coro2a^-/-^*) F_1_ hybrid mice were bred and maintained in the animal facility at our institution, and studies carried out in accordance with the guidelines of our institution’s Animal Care and Use Committee.

### Uterotrophic Assay

Eight week old female mice were ovariectomized (OVX), rested for 1-2 weeks, and treated with E_2_ (40.0 μg/kg BW i.p.) in 0.1 ml saline containing 0.25% ethanol, or ethanol/saline vehicle, at 0, 2, 24, and 48 h. At 24 h after the third treatment (72 h), mice were euthanized and uteri were collected. The uteri were then cleaned of fat and adventitia, blotted to remove luminal fluid, and body weight adjusted uterine wet weights were determined. Uteri were either snap frozen or fixed in 10% formalin.

### Real-time quantitative RT-PCR

RNA was isolated from 1-2 week rested OVX mice at 2 hr following vehicle- and E_2_-treatment. cDNA was then prepared, and *Coro2a* expression analyzed by SYBR Green real-time PCR using methods and primers (forward, GCAATGGAAGCAGTACAAAGCT, reverse, GGTGTGTGCAGATACCAGCG) as previously described (12). Relative RNA equivalents for each sample were obtained by standardization of ribosomal protein L8 levels.

### Immunohistochemical staining

Quantification of uterine eosinophilic infiltrates was carried out as previously described (30). Briefly, frozen sections (6 μm) were cut, air dried for 4-5 h, fixed in acetone for 10 min at −20° C, and then washed in phosphate-buffered saline. Endogenous peroxidase staining of eosinophils was carried out by direct straining with 3,3’-diaminobezidine in PBS with 0.1% H_2_0_2_. Tissue sections were counterstained with Harris’ hematoxylin and mounted.

The slides were examined with an Olympus BX50 upright light microscope (Olympus America, Lake Success, N.Y., USA) with a UPLANApo 20X objective lens via transmitted light. Images were acquired in 24-bit mode with an attached QImaging Retiga 2000R digital ccd camera (Burnaby, BC, Canada). An RGB liquid crystal filter (Cambridge Research Instruments, Woburn MA) was flashed before the camera to create a color image.

At 10× magnification, a grid of premeasured boxes (350 megapixels x 350 megapixels) was laid over each image using Adobe PhotoShop (Adobe Systems, San Jose, California, USA). Boxes within the grid were then numbered sequentially, and an online random number generator was used to select a particular box for quantification (https://www.random.org). The average number of positive cells from five premeasured fields was determined from each sample. In this study, only cross sections of the uteri were used and only fields located entirely within the uterine stroma were quantified.

CORO2A expression was quantified using anti-CORO2A (M-105) rabbit anti-mouse polyclonal antibody (Santa Cruz Biotechnology, Inc., Santa Cruz, CA). Paraffin embedded sections (4 μm) were cut, and mounted onto salinized slides. Slides were deparaffinized using a series of xylene/ethanol washes, followed by a final rinse in dH_2_O. Antigen retrieval was performed using Dako Target Retrieval Solution (Dako, Carpinteria, CA, USA). Slides were incubated in retrieval solution for 20 min, allowed to cool for 20 min, and then washed twice in dH_2_O.

Briefly, the immunohistochemistry protocol was as follows: a 10-min incubation in PBS and 1% BSA, followed by a 15-min incubation in 1% BSA/0.1% Triton X-100 solution. Slides were then washed twice for 10 min/wash in PBS and 1% BSA. Blocking was performed by incubating slides for 30 min in 10% normal goat serum in PBS and 1% BSA. Slides were incubated overnight at room temperature with primary antibody, diluted 1:100 in PBS and 1% BSA. The following day, slides were washed twice for 5 min/wash in PBS and 1% BSA, followed by a 60-min incubation with secondary antibody (goat anti-rabbit IgG Alexa Fluor 555; Invitrogen) diluted 1:500 in PBS and 1% BSA, followed by another 3 washes of 5 min/wash in PBS and 1% BSA. Slides were then incubated for 15 min with DAPI diluted 1:200 in PBS and 1% BSA, washed 3 times for 5 min in PBS and 1% BSA, and coverslips were applied using Aqua Polymount (Polysciences, Inc., Warrington, PA, USA).

Optical sections (2 μm thick) were acquired with a Zeiss 510 META confocal microscope (Carl Zeiss MicroImaging, LLC, Thornwood, NY, USA) using a 40x Plan-Neofluar Apochromat lens. Images were captured using a sequential scan in channel mode with a 1024 × 1024 frame size and 12-bit depth. Signal for CORO2A was captured using a 543 helium neon laser and a long-pass LP 560 emission filter.

Quantification of CORO2A expression was performed using MetaMorph 7.7.3 64-bit offline software (Molecular Devices, Sunnyvale, CA, USA). At least 2 fields/sample were imaged and quantified, and all samples were coded to prevent observer bias. For all samples, an inclusive threshold for intensity was used with a lower limit of 450 and an upper limit of 4095 (default maximum). The regional measurements tool was used to quantify the integrated intensity of CORO2A staining in the luminal epithelium and uterine stroma.

### Statistical Analysis

Statistical analyses were performed using GraphPad Prism version 6.07 and SAS version 9.4 (SAS Institute). The specific tests used to assess the significance of the observed differences are detailed in the figure legends. A P value of ≤0.05 was considered to indicate statistical significance.

### Single Nucleotide Polymorphism and Indel Mapping and Visualization

Single nucleotide polymorphisms (SNPs) and indels between C3H/HeJ and C57BL/6j mice in the region of 4:46535616-46611616 were downloaded from the strain table from Ensembl (http://useast.ensembl.org). The data was parsed and reassembled into BED formatted files in R(3.6.3) for mapping against databases of known regulatory elements binding regions. The Open Regulatory Annotation database (ORegAnno) data was downloaded from the UCSC GenomeBrowser Resource (http://genome.ucsc.edu). This data was also reassembled into a BED file and the intersect function of BEDtools(2.29.2) was used to map the SNPs/indels to the ORegAnno regulatory regions. The ENCODE Candidate Cis-Regulatory Elements (cCREs) was used as a second source of regulatory regions and the data was also collected from the UCSC Genome Browser repository in a bigBed format. The bigBed file was converted into a BED file using the UCSC bigBedtoBED tool(v366) and the intersection of SNPs/indels to the ENCODE cCREs was also conducted using BEDtools. The Integrative Genomics Viewer(2.8.6) (IGV) was used to visualize the SNPs/Indels with the annotated regulatory regions from ORegAnno and ENCODE cCREs in the CORO2A region.

## Results

### Basal uterine epithelial and stromal CORO2A expression co-segregates with differential uterine responsiveness to E_2_

Because we previously identified *Coro2a* as a potential positional candidate for an eQTG underlying *Estq1* (12) (**Fig. 1A**) (F=19.5; P<0.0001), we conducted experiments to localize and quantify the protein level expression of CORO2A in the mouse uterus. In uterine sections, both the epithelium and stroma exhibited positive staining for CORO2A (**Fig. 1B**). Quantification of cellular staining revealed that the epithelial and stromal CORO2A signal differed significantly among the strains as a function of E_2_-treatment. Under basal conditions, CORO2A staining was significantly greater in the epithelium (**Fig. 1C**) (Interaction, P=0.0007; treatment, P<0.0001; strain, P<0.0001) and stroma (interaction, P=0.01; treatment P<0.0001; strain, P<0.0001) (**Fig. 1D**) of high-responder B6 mice compared to low-responder C3H mice. B6C3 mice exhibited intermediate staining, consistent with a haploinsufficiency model (31). In contrast, CORO2A expression was significantly reduced in both the epithelium and stroma at 2 h following treatment with E_2_, and did not differ significantly among the strains.

**Figure 1.**
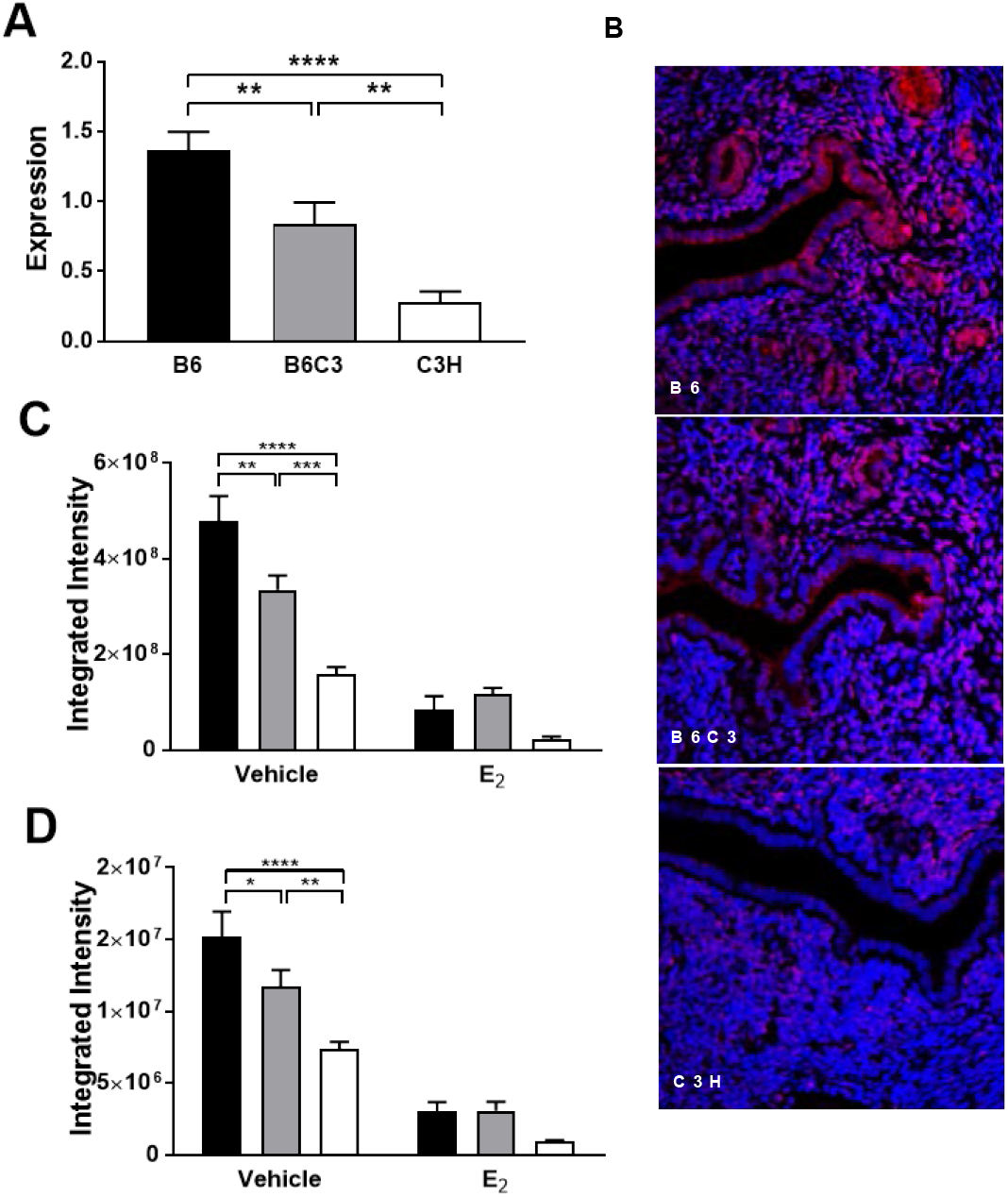
*Coro2a*/CORO2A expression in mouse uterus. Eight week old female B6, B6C3, and C3H mice were ovariectomized (OVX), rested for 1-2 week, treated with E_2_ (40.0 μg/kg BW i.p.) in 0.1 ml saline containing 0.25% ethanol, or ethanol/saline vehicle, and tissues collected at 2 hr. **(A)** Basal uterine *Coro2a* expression differs significantly among vehicle treated OVX mice (F=19.5; P<0.0001). (**B**) Localization of CORO2A in mouse uterus (red, CORO2A; blue, DAPI; ×20 view). Quantity of CORO2A in the uterine (**C**) epithelium and (**D**) stroma differed significantly among the strains (Interaction, P=0.0007; treatment, P<0.0001; strain, P<0.0001 and interaction, P=0.01; treatment P<0.0001; strain, P<0.0001, respectively). Data shown represent the combined results of two experimental cohorts (n=6-16 mice). Significance of observed differences in *Coro2a* and CORO2A expression was determined by one-ANOVA and two-way ANOVA, respectively followed by Holm-Sidak’s multiple comparisons test. *, P<0.05; **, P<0.01; ***, P<0.001; ****, P<0.0001.

### Congenic mapping confirms that *Coro2a* is a positional candidate for the eQTG underlying *Estq1*

The identification of *Coro2a* as a potential positional candidate for an eQTG underlying *Estq1* is based on integrating the 95% confidence interval for linkage of *Estq1* (34.5-59.4 Mbp) to *Coro2a* (46.53-46.60 Mbp) with the finding that genetic control of differential *Coro2a*/CORO2A expression co-segregates with high and low uterine responsiveness to E_2_ (12). In order to confirm the genetic association between *Coro2a* and *Estq1*, we studied the uterine responsiveness of B6.C3H-*Bbaa1a* mice. This congenic mouse line has a C3H-derived interval from 11.6-77.8 Mbp on the B6 background (28, 29) which encompasses both *Estq1* and *Coro2a* (**Fig. 2A**). Following E_2_ treatment, both uterine wet weight (**Fig. 2B**) and the number of stromal eosinophils (**Fig. 2C**) differed significantly among OVX B6, C3H, and B6.C3H-*Bbaa1a* mice (F=9.5; P=0.0008 and F=11.3; P=0.0002, respectively). With respect to uterine wet weight, B6.C3H-*Bbaa1a* mice showed a lower response compared to B6, comparable to C3H. Similarly, the number of stromal eosinophils was significantly lower in B6.C3H-*Bbaa1a* compared with B6, with C3H = B6.C3H-*Bbaa1a* mice. These results demonstrate that the C3H-derived interval encompassing *Estq1/Coro2a* is sufficient to modulate the uterotrophic response, and thus further supports *Coro2a* as a positional candidate for an eQTG underlying *Estq1*.

**Figure 2.**
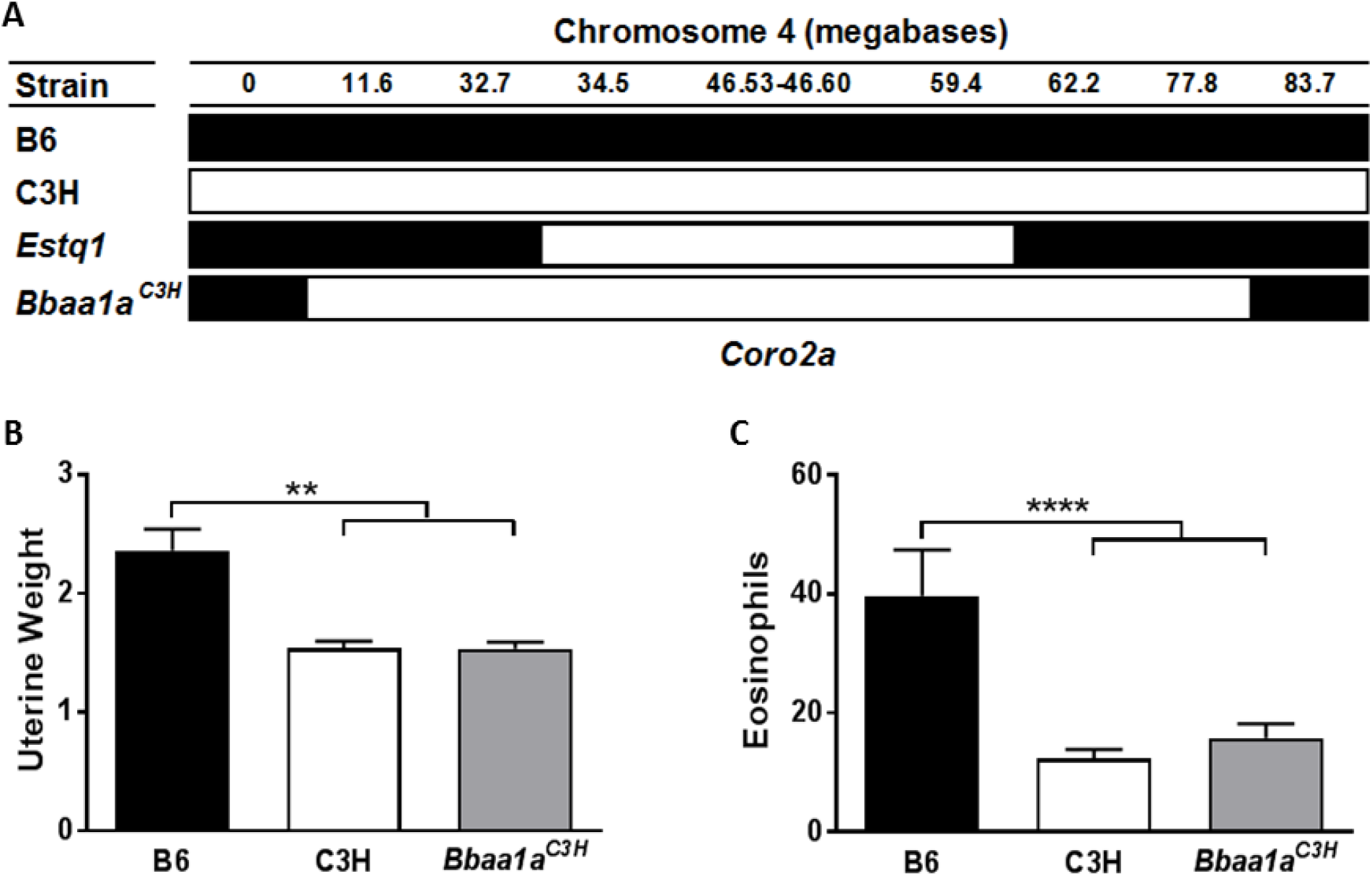
Uterine responsiveness of OVX B6, C3H, and B6.C3H-*Bbaa1a* congenic mice to E_2_. (**A**) Location of *Estq1* and *Coro2a* in B6.C3H-*Bbaa1a* mice. Marker and gene locations are from NCBI annotation of GRCm38.p6. Eight week old female B6, C3H, and B6.C3H-*Bbaa1a* mice were ovariectomized (OVX), rested for 1-2 week, and treated with E_2_ (40.0 μg/kg BW i.p.) in 0.1 ml saline containing 0.25% ethanol, at 0, 24, and 48 h and samples collected at 24 h after the third treatment (72 h). **(B)** Body weight adjusted uterine wet weights (mg/gr) and (**C**) number of eosinophils infiltrating the stroma differed significantly among the strains (F=9.5; P=0.0008 and F=11.3: P=0.0002). Data shown represent the combined results of two experimental cohorts (n=7-14 mice). Significance of observed differences was determined by one-way ANOVA followed by Holm-Sidak’s multiple comparisons test. **, P<0.01; ***, P<0.001; ****, P<0.0001.

### Deficiency-complementation testing establishes *Coro2a* as the eQTG underlying *Estq1*

Given that the differences in uterine wet weight and the number of infiltrating eosinophils co-segregate with *Coro2A*/CORO2A expression among B6, C3H, and B6.C3H-*Bbaa1a* mice, and haploinsufficiency in B6C3 mice, we used B6-*Coro2a*^-/-^ mice and deficiency-complementation testing (26, 27) to examine the causal relationship between *Coro2a* and *Estq1*. Following E_2_-treatment, both uterine weight (**Fig. 3A**) and the number of eosinophils infiltrating the stroma (**Fig. 3B, C**) differed significantly among OVX B6, B6-*Coro2a^B6/-^*, and B6-*Coro2a^-/-^* mice (F=9.2; P=0.0007 and F=27.4; P<0.0001, respectively). The finding that B6-*Coro2a^-/-^* and B6-*Coro2a^B6/-^* mice phenocopy the uterine responsiveness of mice with *Coro2a^C3H/C3H^* and *Coro2a^B6/C3H^* alleles, respectively, indicates that *Coro2a* is the eQTG underlying *Estq1*.

**Figure 3.**
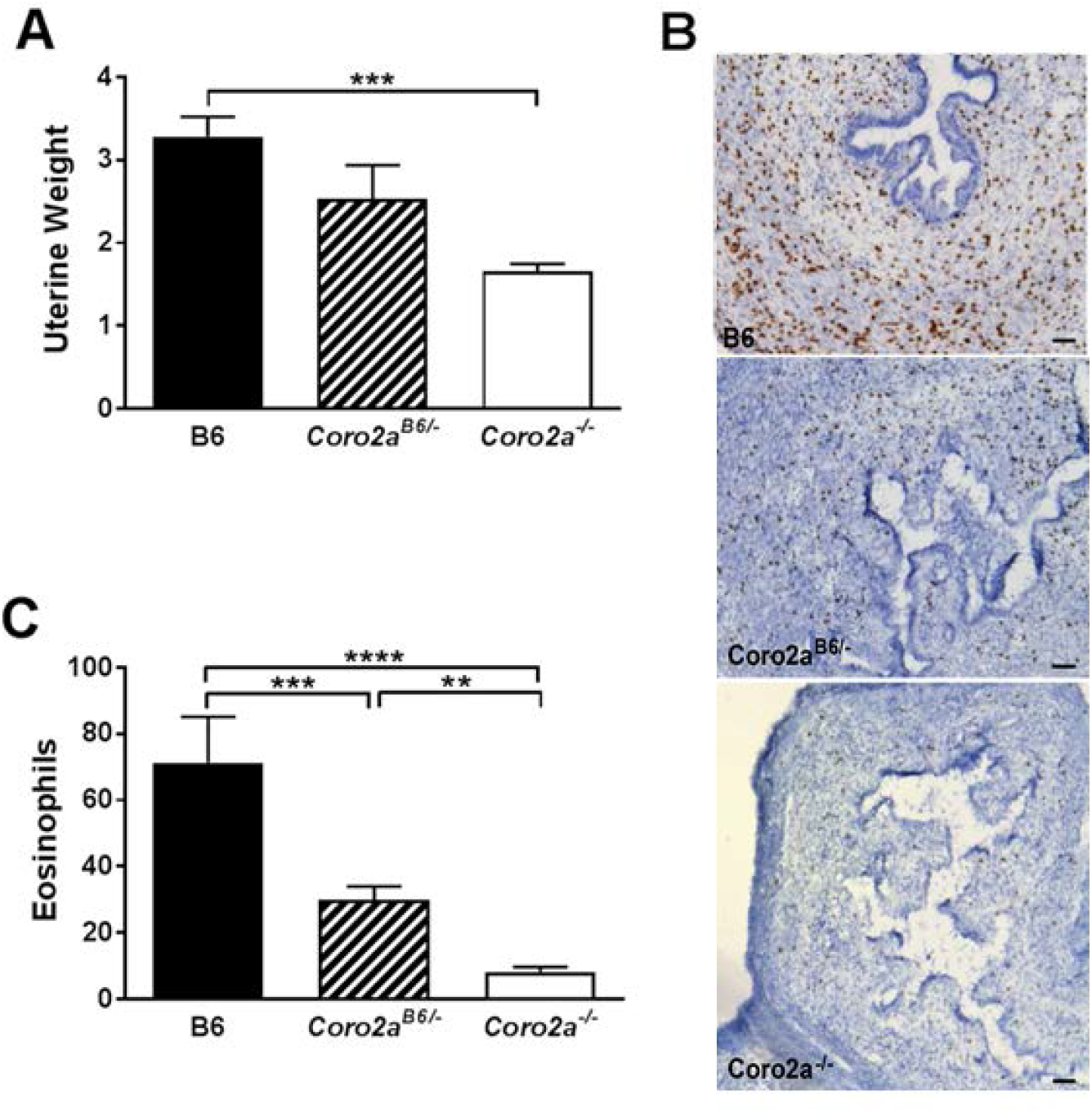
Uterine responsiveness of OVX B6, B6-*Coro2a^B6/-^*, and B6-*Coro2a^-/-^* mice to E_2_. Eight week old female B6, B6-*Coro2a^B6/-^*, and *B6-Coro2a^-/-^* mice were ovariectomized (OVX), rested for 1-2 week, and treated with E_2_ (40.0 μg/kg BW i.p.) in 0.1 ml saline containing 0.25% ethanol, at 0, 24, and 48 h and samples collected at 24 h after the third treatment (72 h). (**A**) Body weight adjusted uterine wet weights (mg/gr) differed significantly among the strains (F=9.2; P=0.0007). (**B**) Endogenous peroxidase staining of eosinophils (brown) in the uterine stroma (blue). (**C**) Average number of eosinophils counted in five premeasured, randomly selected fields differed significantly among the strains (F=27.4; P<0.0001). Data shown represent the combined results of two experimental cohorts (n=7-17). Significance of observed differences was determined by one-way ANOVA followed by Holm-Sidak’s multiple comparisons test. **, P<0.01; ***, P<0.001; ****, P<0.0001.

## Discussion

Taken together, the genetic studies reported herein identify *Coro2a* as the eQTG underlying *Estq1*. To our knowledge, this is the first study to link *Coro2a*/CORO2A directly to E_2_-regulated responses *in vivo*. At baseline, *Coro2a* is expressed at significantly higher levels in the uteri of high responder B6 mice compared to low responder C3H mice (12). We posit that these differences in *Coro2a* expression are due to genetic variation within the *Coro2a* locus regulating transcription. Alignment of strain-specific single nucleotide polymorphisms (SNPs) and indels with experimentally determined cis-regulatory regions and transcription factor binding sites from the ORegAnno database (32) and the mouse ENCODE project (33) identified multiple variations which may mechanistically affect enhancer activity and regulatory factor binding (Supplementary Figure 1 and Tables 1 and 2).

Our study confirmed this differential expression at the protein level, with B6 mice expressing significantly higher levels of CORO2A than low responder C3H mice in both the uterine stroma and epithelium. B6C3 mice, on the other hand, exhibited intermediate levels of CORO2A staining. At 2 hr following E_2_ treatment, CORO2A staining was diminished in both the uterine epithelium and stroma with no significant difference being appreciated between these strains. Together, these data suggest that CORO2A affects differential uterine responsiveness to E_2_ in a concentration dependent manner, doing so at the time of nuclear estrogen receptor (ER) occupancy during the acute/rapid phase (0-1 hr), and not during the late phase (2-24 hr), of the early uterotophic response (34).

Using B6.C3H-*Bbaa1a* mice we confirmed the genetic linkage of *Estq1* to *Coro2a*. Following treatment with E_2_, the uteri of OVX B6 mice exhibited higher body-weight adjusted uterine wet weights and higher levels of stromal eosinophil infiltration when compared to B6.C3H-*Bbaa1a* and C3H mice. Next, deficiency-complementation testing using B6-*Coro2a^-/-^* mice allowed us to assess directly the causal relationship between *Coro2a* and *Estq1*. Here, OVX B6 mice were found to exhibit higher body-weight adjusted uterine wet weights and higher levels of stromal eosinophil infiltration when compared to B6-*Coro2a^B6/-^* and B6-*Coro2a^-/-^* mice following E_2_ treatment.

The finding that the uterine responsiveness of OVX B6-*Coro2a^-/-^* and B6-*Coro2a^B6/-^* mice phenocopy that of C3H and B6.C3H-*Bbaa1a* mice, and B6C3 mice respectively, further supports our hypothesis. In essence, lack of *Coro2a*/CORO2A expression in *Coro2a^-/-^* mice is functionally equivalent to the level of expression associated with C3H mice, which have two copies of the hypomorphic *Coro2a^C3H^* allele. This is further supported by the finding that uterine wet weight and the number of eosinophils infiltrating the stroma of *B6-Coro2a^B6/-^* and B6C3 mice both exhibit haploinsufficiency, where expression of a single copy of the high-responder B6 allele fails to produce enough *Coro2a*/CORO2A to preserve and/or restore high responsiveness (31). Thus, B6 mice require two copies of a functional, normal B6 *Coro2a* gene in order to reach fully functional expression levels. Taken together, our data demonstrate that *Coro2a* is a key eQTG underlying *Estq1*, and raise the possibility that *CORO2A* may be an eQTG contributing to other E_2_-regulated responses and human disease susceptibility.

In this regard, a microarray analysis of differentially expressed genes in invasive breast adenocarcinoma samples identified *CORO2A* as one of the genes that was overexpressed when compared to normal breast tissue (23). In the same dataset, the canonical ER target gene *pS2/TFF_1_* was also overexpressed in tumor samples. A subsequent study also reported that *CORO2A* is overexpressed in malignant breast tissues compared to normal breast tissues (24). *In vitro* functional studies showed that knockdown of *CORO2A* expression inhibited cell migration, decreased viability and colony formation and induced cell cycle arrest in the G_0_/G_1_ phase in breast cancer cells (25). Importantly, in the same study it was shown that patients with elevated expression of *CORO2A* had poorer overall survival. Similarly, increased *CORO2A* expression is associated with shorter progression-free survival times in patients with endometrioid carcinoma and clear cell carcinoma (35), and the malignant phenotype of human colon carcinoma (36). Moreover, *CORO2A rs1985859* has been identified as a GWAS-candidate gene that is predictive of radiosensitivity to preoperative chemoradiotherapy in rectal cancer patients (37), and that downregulation of CORO2A was significantly associated with reduced early apoptosis in colorectal cancer cells *in vitro*. Clearly, *Coro2a*/CORO2A plays a critical role in the genetic control of the murine uterotophic response, and may be a novel oncogene in the initiation and progression of human cancers. Whether or not the observed phenotypes are solely due to CORO2A functioning as an NCoR exchange factor leading to the derepression of nuclear receptor target genes, including ESR1 and ESR2, awaits further experimentation.

## Acknowledgements

The staff at the Microscopy Imaging Center at the University of Vermont, funded, in part, by the National Center for Research Resources (1S10RR019246), for performing staining and imaging of CORO2A. Dr. Christopher K. Glass at the University of California San Diego for providing the B6-^*Coro2atm1(KOMP)Wtsi*^ (B6-*Coro2a^-/-^*) mice. Dr. Janis J. Weis at the University of Utah for providing the B6.C3H-*Bbaa1a* (11.6-77.8 Mbp) mice.

## Author Contributions

J. Findley performed research; C. Teuscher designed research. J. Findley and C. Teuscher analyzed data. J. Findley, E. McGee, C.-Y. Lin, D. Krementsov and C. Teuscher wrote and edited the paper.

