## Supplemental Figure 1 for "*Coro2a*, An Expression Quantitative Trait Gene Underlying *Estq1*, Controls Uterine Responsiveness to Estradiol"

### Slide 1
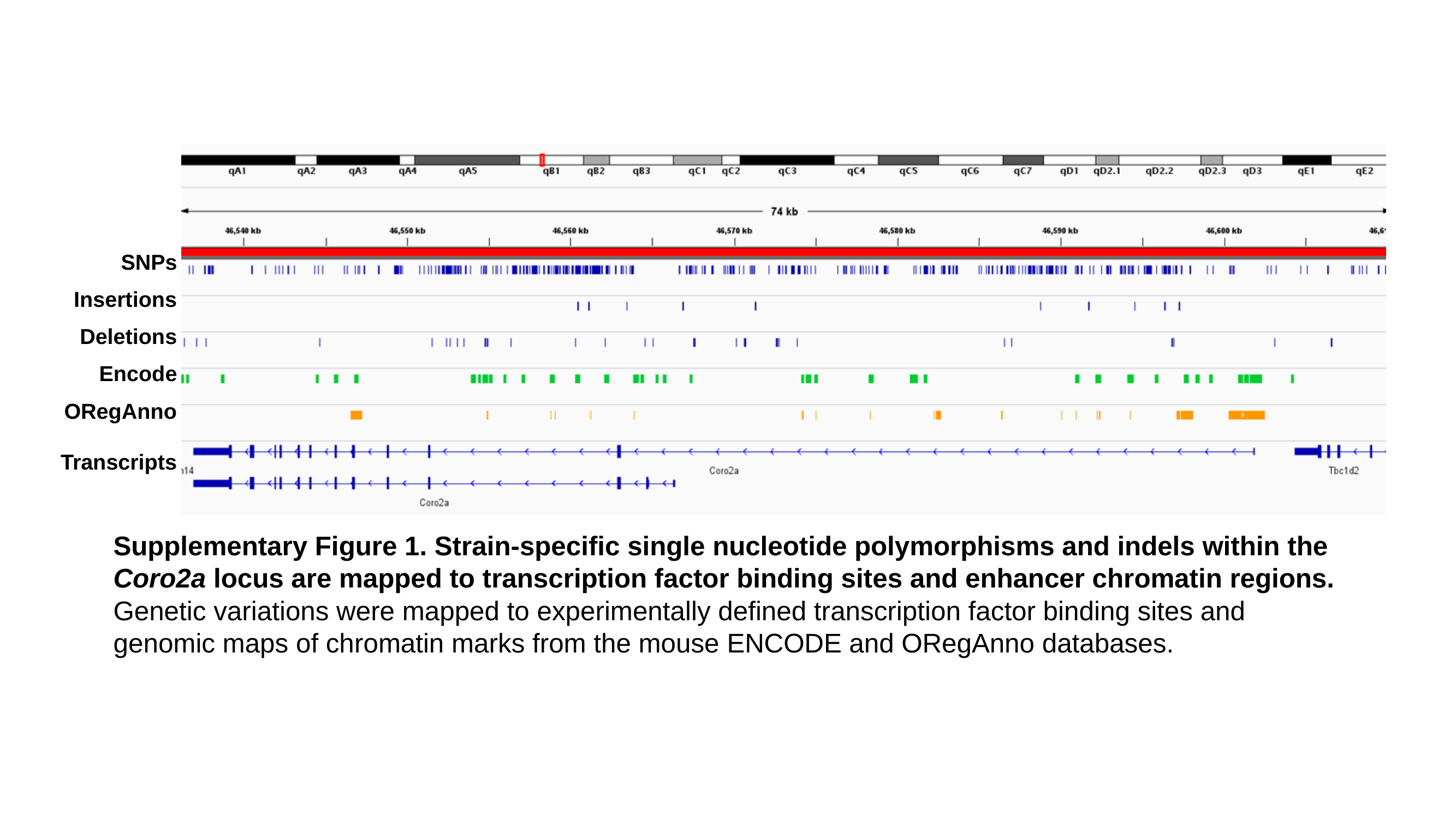

SNPs
Insertions
Deletions
Encode
ORegAnno
Transcripts
Supplementary Figure 1. Strain-specific single nucleotide polymorphisms and indels within the Coro2a locus are mapped to transcription factor binding sites and enhancer chromatin regions. Genetic variations were mapped to experimentally defined transcription factor binding sites and genomic maps of chromatin marks from the mouse ENCODE and ORegAnno databases.
